# ReactomeGSA - Efficient Multi-Omics Comparative Pathway Analysis

**DOI:** 10.1101/2020.04.16.044958

**Authors:** Johannes Griss, Guilherme Viteri, Konstantinos Sidiropoulos, Vy Nguyen, Antonio Fabregat, Henning Hermjakob

## Abstract

Pathway analyses are key methods to analyse ‘omics experiments. Nevertheless, integrating data from different ‘omics technologies and different species still requires considerable bioinformatics knowledge.

Here we present the novel ReactomeGSA resource for comparative pathway analyses of multi-omics datasets. ReactomeGSA can be used through Reactome’s existing web interface and the novel ReactomeGSA R Bioconductor package with explicit support for scRNA-seq data. Data from different species is automatically mapped to a common pathway space. Public data from ExpressionAtlas and Single Cell ExpressionAtlas can be directly integrated in the analysis. ReactomeGSA thereby greatly reduces the technical barrier for multi-omics, cross-species, comparative pathway analyses.

We used ReactomeGSA to characterise the role of B cells in anti-tumour immunity. We compared B cell rich and poor human cancer samples from five TCGA transcriptomics and two CPTAC proteomics studies. There, B cell-rich lung adenocarcinoma samples lack the otherwise present activation through NFkappaB. This may be linked to the presence of a specific subset of tumour associated IgG+ plasma cells that lack NFkappaB activation in scRNA-seq data from human melanoma. This showcases how ReactomeGSA can derive novel biomedical insights by integrating large multi-omics datasets.

---

Increasingly available approaches such as transcriptome sequencing (RNA-seq), mass spectrometry (MS)-based shotgun proteomics, and microarray studies enable us to characterise genome- and proteome-wide expression changes. This leads to the challenge of deriving relevant biological insights from lists of hundreds of regulated genes and proteins.

Pathway analysis techniques have emerged as a solution to this problem. Resources like the Gene Ontology (GO) ^1^, the Kyoto Encyclopedia of Genes and Genomes (KEGG) ^2^, the Molecular Signatures Database (MSigDB) ^3^, or Reactome ^4^ organise existing biological knowledge into gene sets or pathways. Pathway analysis approaches can use these resources to represent long lists of regulated genes and proteins as biologically defined pathways. This leads to a more intuitive interpretation of the data and increases the statistical power. While single genes or proteins may only show small, non-significant changes, synchronous changes within a pathway may reveal a biologically important effect. Thereby, pathway analysis has become an essential resource for ‘omics data analyses.

The increasing availability of public ‘omics datasets has made it common practise to include these into analyses. This data integration is commonly complicated as datasets were created in different species or using different ‘omics approaches. Pathway analysis approaches offer a solution to this problem since data can be mapped to the more general and comparable pathway space.

Existing web-based pathway analysis resources, such as PANTHER ^5^, the Database for Annotation, visualisation and Integrated Discovery (DAVID) ^6^ or Reactome’s pathway analysis ^7^ all provide over-representation analyses. This type of pathway analysis only tests whether a list of genes is overrepresented in a specific pathway. These approaches have the advantage that the user input is simple, but ignore any underlying quantitative information at the cost of reduced statistical power. Moreover, users have to manually separate up- and down-regulated genes and process them in separate analyses. Thereby, any result is only a partial representation of the underlying biological changes.

The recently developed iLINCS resource extends the concept of single-resource pathway analysis to a powerful multi-omics and multi-resource analysis ^8^. It tests whether a list of gene / protein identifiers correlates with a large set of pre-computed signatures. These signatures are often the result of differential expression analyses. Therefore, similar to the aforementioned resources, iLINCS ignores any underlying quantitative information in the final comparison. Additionally, the comparison with public data is limited to pre-defined experimental designs and comparisons whose results are stored as pre-computed signatures. Therefore, a large portion of the data remains unused.

Here, we present the novel Reactome gene set analysis system “ReactomeGSA”. ReactomeGSA supports the comparative pathway analysis of multiple independent datasets. Datasets are submitted to a single pathway analysis and represented side-by-side on the pathway level. It uses gene set analysis methods that take the quantitative information into consideration and thereby performs the differential expression analysis directly on the pathway level. Data from different species is automatically mapped to a common pathway space through Reactome’s internal mapping system. All supported gene set analysis methods are optimised for different types of ‘omics approaches including single cell RNA-sequencing (scRNA-seq) data. Public datasets can be directly integrated from ExpressionAtlas and Single Cell ExpressionAtlas ^9^. We used ReactomeGSA to show that B cell receptor signalling is surprisingly down-regulated in B cell-rich lung adenocarcinoma in contrast to four other human cancers. We could further link this to IgG+ plasma cells in scRNA-seq data. ReactomeGSA thereby provides easy access to multi-omics, cross-species, comparative pathway analysis to reveal key biological mechanisms by integrating large ‘omics datasets.

## Results

ReactomeGSA can be accessed through Reactome’s web interface (https://www.reactome.org/PathwayBrowser/#TOOL=AT) or through the novel ReactomeGSA R Bioconductor package (https://doi.org/doi:10.18129/B9.bioc.ReactomeGSA, Figure 1). Both access the public application programming interface (API) to perform the pathway analysis. The analysis system is a Kubernetes application based on the microservice paradigm that automatically scales to current demand (see Methods for details). This infrastructure enables us to offer computationally expensive pathway analysis methods through an open interface. ReactomeGSA currently supports three methods: PADOG ^10^, Camera through the limma R package ^11^, and the single-sample gene set enrichment analysis (ssGSEA) ^12^ through the GSVA ^13^ R package (see Methods for details). The API and its complete specification is publicly available at https://gsa.reactome.org. Thereby, ReactomeGSA can easily be integrated into any other software infrastructure.

**Figure 1.**
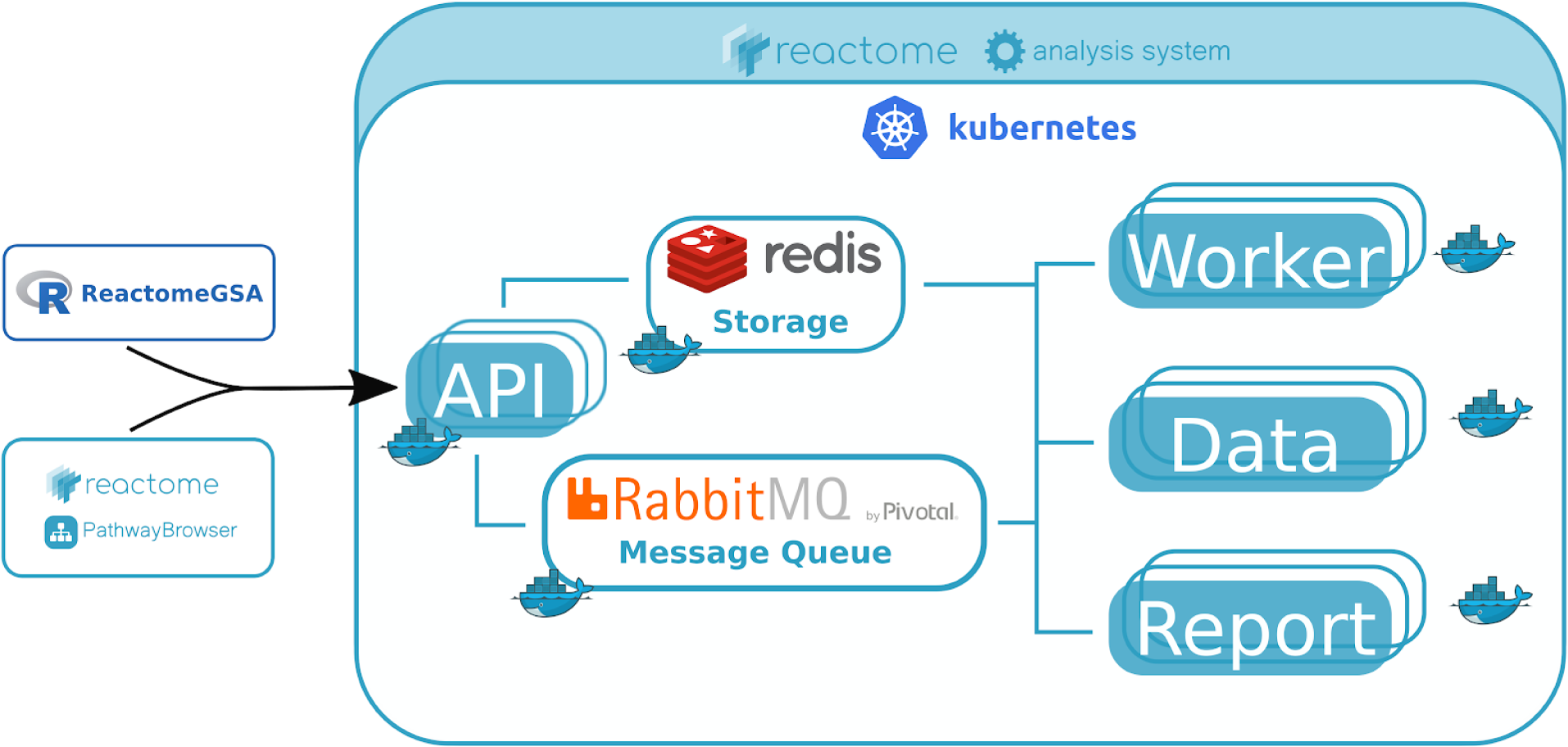
Schema of the ReactomeGSA system. All requests are sent to a public web-based API through the ReactomeGSA Bioconductor R package or Reactome’s web-based PathwayBrowser. The system is a Kubernetes application based on the microservices architecture. All requests are distributed through an internal message queue using RabbitMQ. Worker nodes are responsible for the complete pathway analysis, including identifier mapping and the creation of the visualisation data in Reactome’s pathway browser. Data nodes are responsible to load data from external resources such as ExpressionAtlas. Finally, report nodes create PDF and Microsoft Excel files as a static report of the analysis results. All data is stored in a central Redis instance. All nodes are Docker containers that are orchestrated by Kubernetes and automatically scaled based on current demand. Thereby, the application can dynamically adapt to changing usage levels.

ReactomeGSA is fully integrated in Reactome’s existing web-based pathway browser application (Figure 2). After choosing the new “Analyse gene expression” tab and the desired analysis method, the user can add any number of datasets to the analysis request. Public datasets are directly loaded from Expression Atlas and the Single Cell Expression Atlas ^9^. Results can be sent as emails including static PDF and Microsoft Excel reports. Finally, the complete gene set analysis result is visualised in Reactome’s interactive pathway browser. The pathway browser enables users to view Reactome’s complete pathways from a tree-based, hierarchical overview, down to the single gene- and protein-level reactions. The results of different datasets can be switched at the click of a button or automatically changed every few seconds like a slideshow across all results. Thereby, differences between the analysed datasets are immediately visible and can subsequently be interactively investigated down to the single gene or protein level.

**Figure 2.**
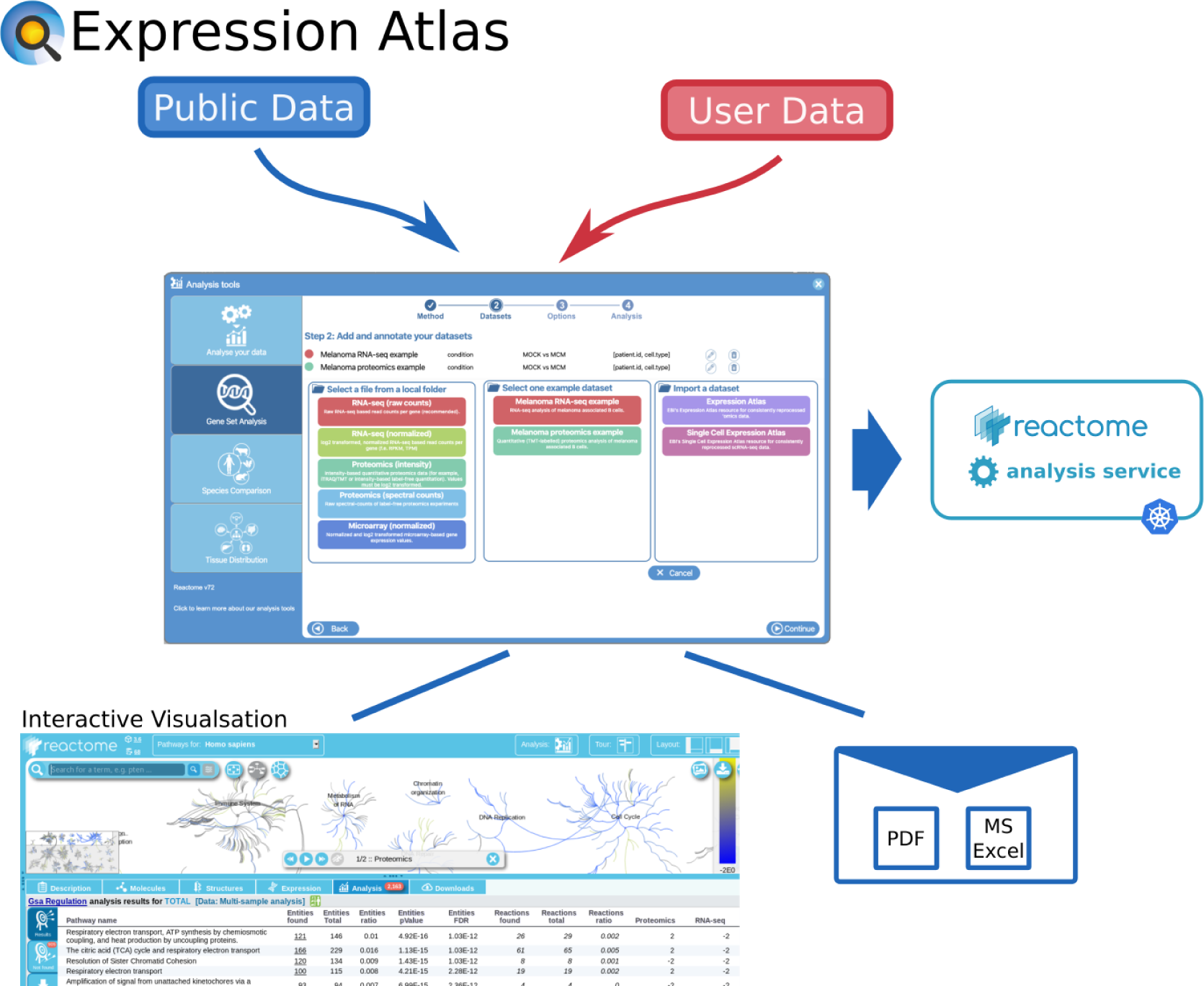
ReactomeGSA is fully integrated into the web-based Reactome pathway browser (https://reactome.org). Users can either upload their own datasets or import public data from ExpressionAtlas. The gene set analysis is performed through the ReactomeGSA API. Results are visualised in Reactome’s interactive pathway browser and send as static reports in PDF and Microsoft Excel format via email.

The ReactomeGSA R package has been included in Bioconductor since version 3.10 (Figure 3). Similar to the web interface, multiple datasets can be added to a ReactomeAnalysisRequest object. Expression values and metadata can directly be loaded from Bioconductor ExpressionSet, limma EList ^11^ and edgeR ^14^ DGEList objects. Thereby, the ReactomeGSA package can easily be integrated into existing R-based workflows. The analysis results are returned as a ReactomeAnalysisResult object. This object contains the pathway analysis results across all analysed datasets, as well as the gene- or protein-level results of the differential expression analysis. It can directly open the interactive visualisation in Reactome’s web-based pathway browser (see above) and create plots to visualise the comparative pathway analysis results. Thereby, the multi-dataset results generated by ReactomeGSA can be natively processed in R.

**Figure 3.**
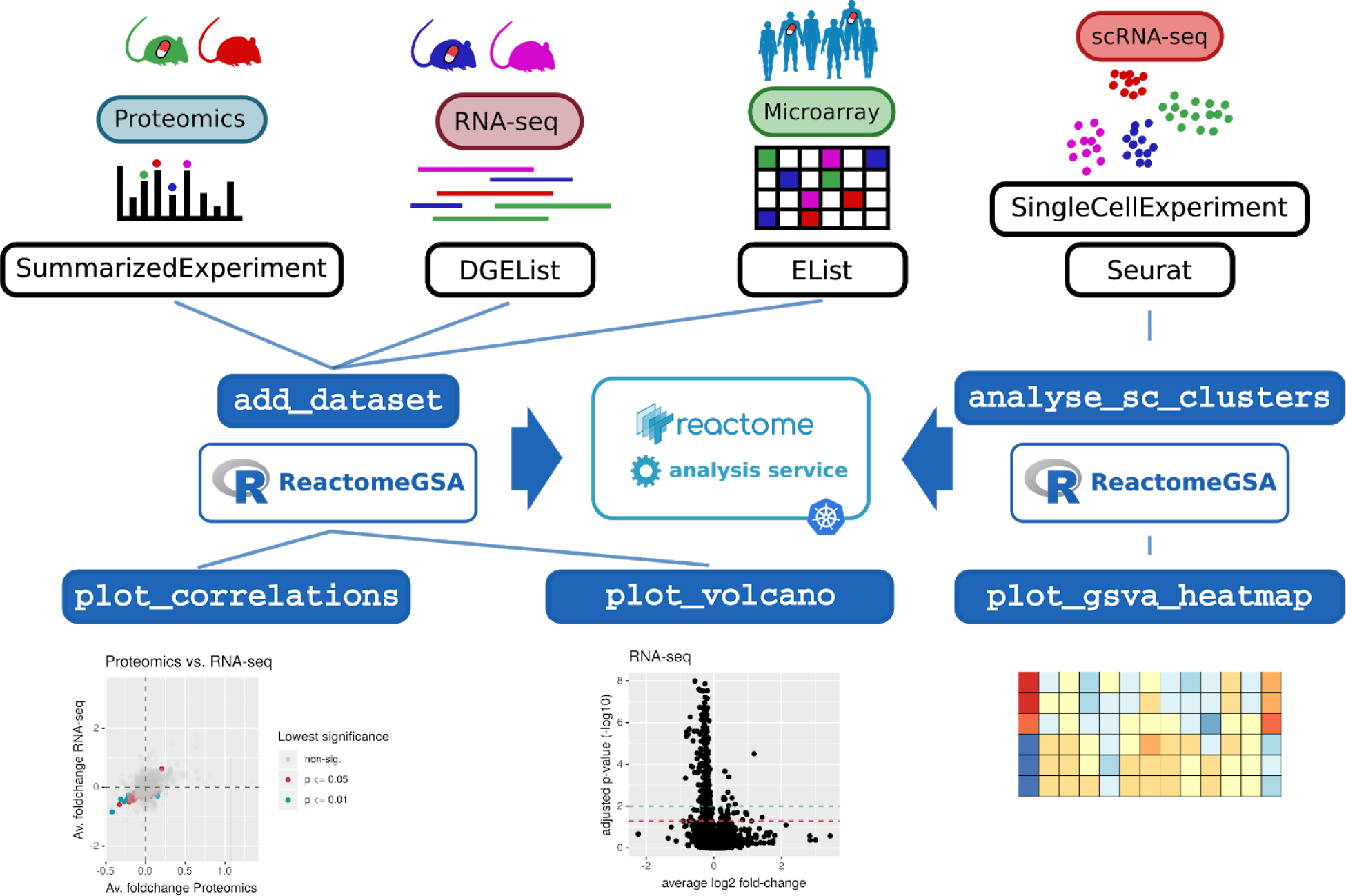
The ReactomeGSA Bioconductor R package can directly process data from the most commonly used data structures for ‘omics analyses. The pathway analysis is performed through the ReactomeGSA analysis system and made available through a native R object. Convenient plotting functions give a quick overview of how well two datasets correlate on the pathway level. Volcano plots further highlight the magnitude of the observed changes in individual datasets. Additionally, pathway analysis of scRNA-seq data is simplified through the single “analyse_sc_clusters” function.

The ReactomeGSA R package has dedicated features to simplify pathway analyses of scRNA-seq data (Figure 3). The “analyse_sc_clusters” function can directly process Seurat ^15^ and Bioconductor’s SingleCellExperiment objects ^16^. It automatically retrieves the average gene expression per cell cluster and performs an ssGSEA analysis on the cluster-level expression values. This results in one pathway-level expression value per cell cluster. Thereby, cell clusters can quickly be interpreted based on specific biological functions.

### ReactomeGSA Reveals a Lack of B Cell Activation in B Cell-rich Lung Adenocarcinoma

We were among the first to show that B cells play a crucial role in anti-tumour immunity in human melanoma ^17^. *In vitro*, B cells differentiate towards a tumor-induced, plasmablast-like (TIPB) phenotype in the presence of melanoma cells. The corresponding molecular TIPB signature predicts overall survival in the The Cancer Genome Atlas (TCGA) melanoma cohort. Whether this effect is specific to melanoma or whether it is a general part of the anti-tumour immune response is currently unknown.

We analysed the difference between TIPB-high *vs.* TIPB-low samples in the TCGA cohorts for melanoma ^18^, lung adenocarcinoma ^19^, lung squamous cell carcinoma ^20^, ovarian cancer ^21^, and breast cancer ^22^. Melanoma and ovarian cancer patients with high levels of TIPB showed significantly longer overall survival (likelihood ratio test p < 0.01 for both, hazard ratio 0.56 melanoma, 0.69 ovarian cancer, Figure 4A). There was no significant difference in overall survival for lung adenocarcinoma, lung squamous cell, and breast cancer patients (likelihood ratio test p = 0.04, p = 0.2 and p = 0.9 respectively). Therefore, the effect of TIPB on anti-tumour immunity and patient survival differs across these types of cancers.

**Figure 4.**
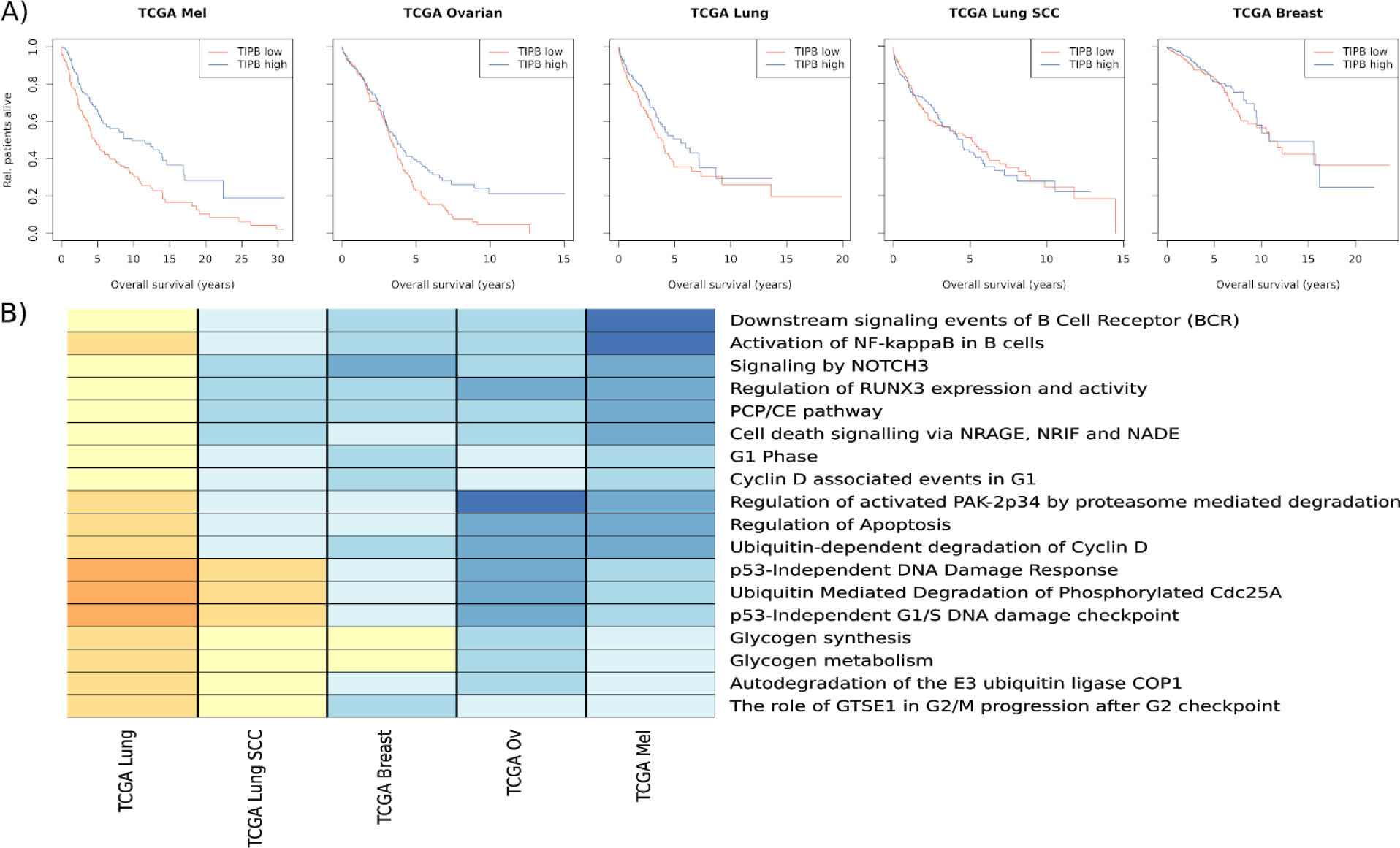
Comparison of TIPB-high vs. -low samples from TCGA studies on melanoma (TCGA Mel), ovarian cancer (TCGA Ovarian), lung adenocarcinoma study (TCGA Lung), lung squamous cell carcinoma (TCGA Lung SCC), and breast cancer (TCGA Breast). **A)** Overall survival of patients with high (blue line) or low (red line) expression of the TIPB signature (split by the median expression in the dataset). **B)** Average gene fold-changes per pathway. Only pathways significantly regulated (FDR < 0.1) in the TCGA melanoma and the TCGA lung adenocarcinoma cohort with a different direction of regulation in these two cohorts are shown. Shades of yellow represent a down-regulation, shades of blue an up regulation.

We subsequently assessed pathway-level differences between patients with high- and low-levels of TIPB in the five cohorts. The comparative pathway analysis was performed using our ReactomeGSA R package and the PADOG gene set enrichment analysis. 383 pathways were significantly regulated in at least one of the datasets (FDR < 0.1, Supplementary Data 1). 64 of these pathways showed a differential regulation in one of the datasets compared to melanoma. We previously showed *in vitro* that NF-kappaB activation was significantly up-regulated in B cells after stimulation with melanoma conditioned medium ^17^. Lung adenocarcinoma samples were the only ones that showed a significant down-regulation of the “Activation of NF-kappaB in B cells” pathway (FDR = 0.08). Even though these samples have a higher number of TIPB, overall B cell activation is reduced.

We, therefore, assessed how the lung adenocarcinoma cohort differs from the melanoma cohort. In total, 18 pathways were significantly regulated in both the melanoma and the lung adenocarcinoma cohort (Figure 4B). Next to the down regulation of NF-kappaB related genes, there was an overall down-regulation of B cell receptor signalling, but also p53 related DNA damage response, cell cycle and apoptosis related pathways. This shows that lung adenocarcinoma samples with a high number of tumor induced plasmablast-like B cells have a distinct different signalling state compared to melanoma.

Pathways related to B cell receptor signalling and apoptosis correlate with the survival benefit observed through higher numbers of TIPB. The melanoma and ovarian cancer cohort both showed the strongest survival benefit which was linked to the strongest up-regulation of apoptosis related pathways but also B cell receptor signalling. These results highlight that ReactomeGSA’s comparative pathway analysis can quickly reveal clinically relevant conserved signalling events.

### Cancer-relevant Pathways Differ in Proteomics and Transcriptomics Data

In our recent characterisation of melanoma associated B cells, key phenotypic changes in B cells were primarily observed on the protein but not the transcriptome level. We therefore performed a comparative pathway analysis of the two TCGA cohorts where matched whole-proteome profiling from the Clinical Proteomic Tumor Analysis Consortium (CPTAC) are available.

99 samples of the breast cancer CPTAC study ^23^ and 62 samples of the CPTAC ovarian study ^24^ could be directly mapped to samples from the respective TCGA study. As our TIPB signature was only validated for transcriptomics data, sample grouping into TIPB-high and -low samples was transferred from the TCGA data. The pathway analysis was performed using our ReactomeGSA R package and PADOG. 113 and 96 pathways were significantly regulated (FDR < 0.05, Supplementary Data 2) in the proteomics and transcriptomics data from the breast and ovarian cancer study respectively. Out of these, 13 showed a different direction of regulation in the breast study, and one in the ovarian cancer study. In breast cancer, these included VEGF signalling, EGFR signalling, and IGF1R signalling related pathways (all up-regulated in transcriptomics and down-regulated in proteomics). In ovarian cancer, FGFR signalling was significantly up-regulated in the transcriptomics but down-regulated in proteomics data. All of these pathways are linked to proliferation and are relevant pathways to tumour biology. B cell receptor signalling associated pathways were significantly up-regulated in all datasets. This highlights how ReactomeGSA can quickly reveal biologically relevant differences and similarities between ‘omics datasets.

### IgG+ Plasma cells Show Reduced NFϰB Activation

Specific subtypes of B cells seem to be primarily responsible for the B cell triggered anti-tumour response ^17,25–27^. We therefore assessed whether the observed difference in NFϰB activation is B cell subtype specific.

The extracted B cells from the scRNA-seq dataset by Jerby-Arnon *et al.*^*28*^ formed 13 distinct clusters using Seurat (see Methods for details). Based on canonical B cell markers ^29^ we classified these clusters as double negative B cells, seven types of memory-like B cells, memory-switched resting and -activated B cells, naive B cells, plasma cells, and plasmablast-like B cells (Figure 5A). Consistent with their transitional phenotype between B cells and plasma cells, plasmablast-like B cells were the only to express SDC1 (CD138) and low levels of MS4A1 (CD20). This classification already highlights issues in classifying B cell subtypes as we had to classify seven clusters as memory B cells even though they showed marked differences in overall gene expression.

**Figure 5.**
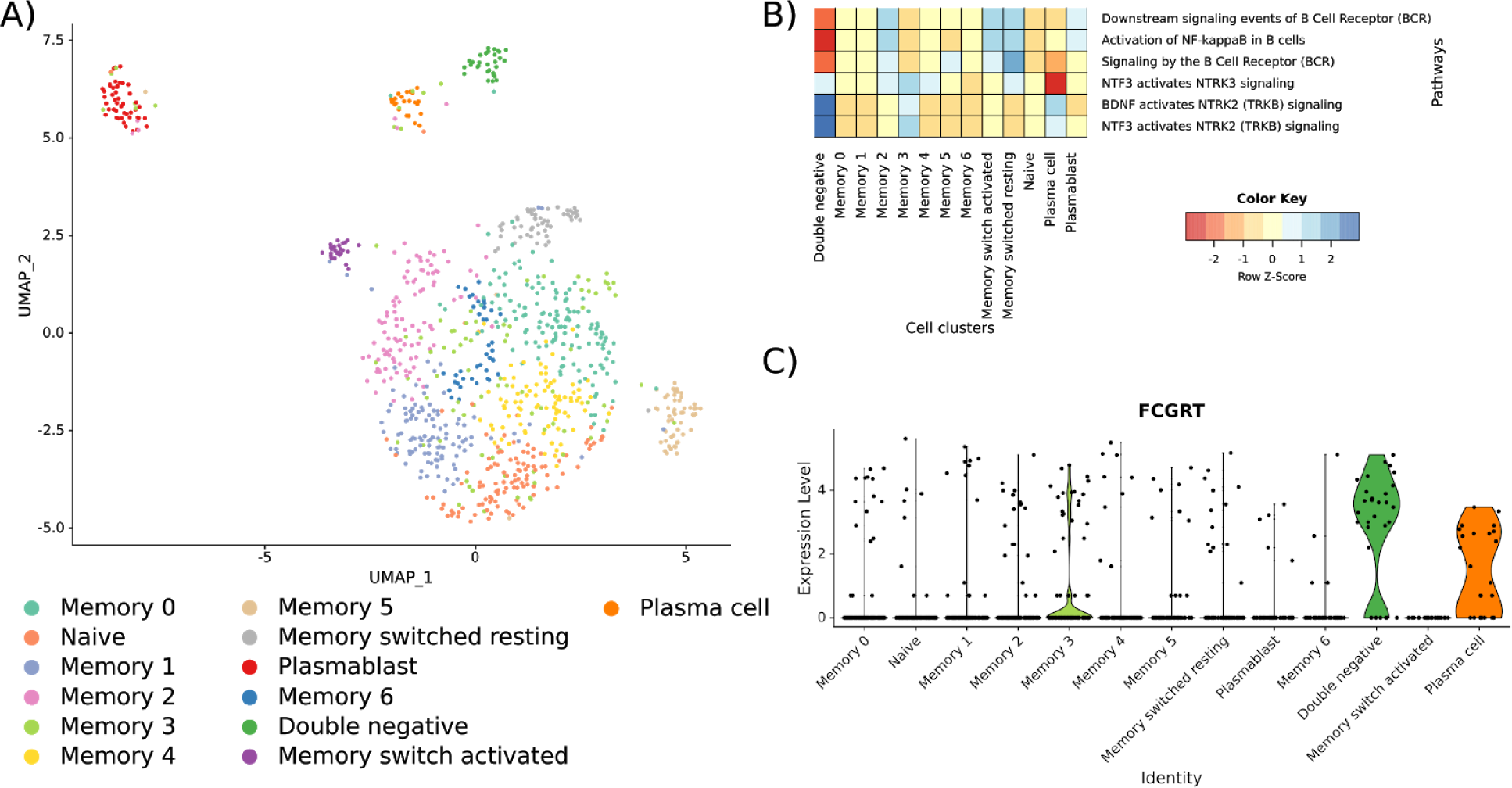
Analysis of B cell subtypes from the dataset by Jerby-Arnon *et al.*^*28*^ **A)** UMAP plot of the identified B cell clusters. Cell type annotations are based on canonical B cell markers ^29^. **B)** ReactomeGSA gene set variation based pathway-level expression in the identified B cell clusters of the Jerby-Arnon *et al.* dataset. Expression values were z-score normalised by pathway. **C)** Expression of IgG estimated through FCGRT abundance in the B cell clusters

We used ReactomeGSA R package’s analyse_sc_clusters function to quantify pathways in these B cell clusters. There was a considerable heterogeneity between the memory B cell clusters, as well as plasmablast and plasma cells in terms of B cell receptor signalling (Figure 5B). In the latter, this matches the previously described lack of functional B cell receptors in IgG positive plasma cells ^30^. Consistently, plasma cells but not plasmablast-like B cells expressed high levels of IgG as determined through Fc fragment of IgG receptor and transporter (FCGRT) expression (Figure 5C). Plasma cells and plasmablast-like B cells further differed in NTRK signalling which regulates cell survival, proliferation and motility ^31^. Our original TIPB signature is too coarse to perfectly differentiate between plasma cells and plasmablast-like B cells. Therefore, the lack of B cell receptor signalling in lung adenocarcinoma samples points towards the high abundance of IgG+ plasma cells. These were shown to be negative prognostic factors in lung adenocarcinoma ^32^ which may explain the reduced survival benefit of TIPB there.

## Discussion

ReactomeGSA greatly decreases the technical challenge to perform pathway analyses of unrelated datasets irrespective of ‘omics technology and investigated species. The iLINCS resource ^8^ is comparable in terms of the integration of different ‘omics data types and public datasets. In contrast to iLINCS, ReactomeGSA does not rely on pre-computed signatures for public datasets. This limits the number of public datasets that can be integrated into a single analysis. At the same time, it gives the researcher complete freedom in terms of experimental design and data analysis strategy to use. Our analysis of TCGA datasets based on a custom signature, for example, would not be supported by iLINCS. Additionally, ReactomeGSA directly supports quantitative ‘omics data as input. Thereby, we can use gene set analysis approaches with increased statistical power compared to simple overrepresentation analysis ^33^. Moreover, the support for sample-level quantitative data enables us to integrate gene set variation analyses which we found especially helpful in the analysis of scRNA-seq data. We, thus, believe that the ReactomeGSA system is a considerable step forward in giving researchers easy access to compex, more sophisticated pathway analysis methods.

A key decision in multi-omics pathway analyses is how to integrate different types of ‘omics data. Methods such as the Gene Set Omic Analysis (GSOA) ^34^ or the PAthway Recognition Algorithm using Data Integration on Genomic Models (PARADIGM) ^35,36^ merge different ‘omics measurements into a single result. Thereby, only data from the same or highly similar samples can be integrated. Moreover, differences between the different ‘omics measurements disappear. As highlighted in our example data and previous studies, such differences are to be expected ^17,23^. We deliberately developed a system that can highlight such differences that researchers can interactively investigate with the Reactome pathway browser. ReactomeGSA therefore provides a novel multi-omics pathway analysis infrastructure that is tailored to expert bioinformaticians and non-experts alike.

### Online Methods

The ReactomeGSA analysis system is accessible through a web-based application programming interface (API). We provide two end-points to this API: an integration into Reactome’s web-based pathway browser application and the ReactomeGSA R Bioconductor package.

The backend is a Kubernetes application (https://kubernetes.io/) currently consisting of six deployments. Each deployment represents one Docker container (Docker Inc, https://www.docker.com). All data is stored in a Redis instance (https://redis.io/). The different components are linked through a message system provided by RabbitMQ (Pivotal, https://www.rabbitmq.com/). All components of the ReactomeGSA backend are developed in Python. The actual gene set analysis is performed using R Bioconductor ^37^ packages through the rpy2 (https://rpy2.github.io/) Python interface to the R language in the worker node (Figure 1).

A key advantage of this setup is that the complete ReactomeGSA application can be described in one so-called YAML file - a Kubernetes configuration file. Since all Docker containers are freely available on Docker Hub (https://hub.docker.com) the ReactomeGSA system can be deployed using the single “kubectl apply -f reactome_gsa.yaml” command. We created a single YAML-formatted configuration file to quickly adapt ReactomeGSA to different use cases (ie. the number of resources available to the different nodes). Detailed information on how to adapt ReactomeGSA can be found on the GitHub repository (https://github.com/reactome/ReactomeGSA). Thereby, users can set up their own version of the ReactomeGSA system within minutes and deploy it locally or in the cloud.

### Multi-omics Gene Set Analysis

At the time of writing, ReactomeGSA supports three different analysis methods: Camera through the limma ^11^ package, PADOG ^10^, and the single-sample gene set enrichment analysis (ssGSEA) ^12^ through the GSVA ^13^ package. All pathway analyses are performed by the worker node in the ReactomeGSA system (Figure 1).

Reactome annotation focuses on human pathways. Thus, as a first step in the analysis, the submitted identifiers are mapped to human UniProt ^38^ identifiers using Reactome’s identifier mapping system. A key issue in mapping identifiers between different identifier systems and across species is to resolve one-to-many mappings. In these cases, the ReactomeGSA system keeps an internal record of these mappings. When mapping the observed genes / proteins to pathways, genes that map to multiple UniProt identifiers which all belong to the same pathway are only added once to this pathway. Thereby, one-to-many mappings are resolved at the pathway-level and inaccuracies normally introduced through identifier conversions are greatly reduced.

In order to increase the coverage of Reactome pathways, pathways can be extended through medium and high confidence interactions derived from IntAct ^39^. This function considerably extends Reactome’s coverage.

At the time of writing, the ReactomeGSA system supports five types of quantitative ‘omics data: Microarray intensities, transcriptomics raw and normalised read counts, and proteomics spectral counts and intensity-based quantitative data. Internally, these different types of data are processed using two different methods: statistics for discrete quantitative data (in case of raw transcriptomics read counts and spectral counting based quantitative proteomics data) and statistics for continuous data. For Camera and PADOG, discrete values are normalised using edgeR’s ^14^ calcNormFactors function. Then, the data is transformed using limma’s voom function ^40^. Continuous data is directly processed using limma ^11^ and normalised using limma’s normalizeBetweenArrays function. The pathway analysis is subsequently performed using limma’s camera function or PADOG as implemented in the respective Bioconductor R package ^33^. For the ssGSEA method ^12^ the analysis is performed using the GSVA Bioconductor R package ^13^. Discrete data is processed using a poisson kernel and continuous data using a gaussian kernel. Thereby, multiple types of ‘omics data can be supported.

### scRNA-seq Pathway Analysis

The analysis of scRNA-seq data is supported through the ReactomeGSA R package’s “analyse_sc_clusters” function, as well as through the direct import of data from the Single Cell Expression Atlas ^9^. In both cases, we calculate the mean expression of genes within a cluster. For the R package, this is done through either Seurat’s ^15^ AverageExpression function, or through scater’s ^41^ “aggregateAccrossCells” function depending on the input object. Single cell data retrieved from the Single Cell Expression Atlas is processed using custom python code (see https://github.com/reactome/gsa-backend for details). This approach to create pseudo-bulk RNA-seq data resembles previously described methods to calculate differentially expressed genes ^16^. Thereby, all pathway analysis methods supported by the ReactomeGSA analysis system are accessible to scRNA-seq data as well.

### TCGA B Cell Analysis

The TCGA transcriptomics data for melanoma (TCGA-SKCM) ^18^, lung adenocarcinoma (TCGA-LUAD) ^19^, lung squamous cell carcinoma (TCGA-LUSC) ^20^, ovarian cancer (TCGA-OV) ^21^, and breast cancer (TCGA-BRCA) ^22^ were retrieved using the TCGAbiolinks R Bioconductor package ^42^. For all datasets apart from melanoma, only primary tumour samples were retained. Genes that were expressed in less than 30% of the samples with at least 10 reads were removed.

The abundance of plasmablast-like B cells (TIPB) was quantified using the single-sample Gene Set Enrichment Analysis (ssGSEA) method ^12^ as implemented in the GSVA R Bioconductor package ^13^. Plasmablast-like B cells were described as CD38, CD27, and PAX5 ^17^. Samples were classified as TIPB-high and -low split by the median expression of the TIPB signature in all samples of the cohort. Overall survival was assessed using the R survival package.

The comparative pathway analysis was performed using the ReactomeGSA R Bioconductor package. In all studies, plasmablast “high” and “low” samples were compared with each other using PADOG ^10^.

The complete R code of this analysis, including the detailed versions of all R packages used is available in the respective Jupyter notebook (see Data availability).

### CPTAC Data Analysis

Data processed through the common data analysis pipeline (CDA) was downloaded from the CPTAC data portal (breast cancer at https://cptac-data-portal.georgetown.edu/cptac/s/S015, ovarian cancer at https://cptac-data-portal.georgetown.edu/cptac/s/S020). For breast cancer ^23^, we used the proteome-level iTRAQ summary, for ovarian cancer ^24^ the PNNL-based protein-level iTRAQ summary. Samples were matched to the respective TCGA samples through the short barcode using the first 11 characters. Only unambiguous matches were retained. Plasmablast abundance based groupings were transferred from the respective TCGA dataset. The data was analysed using the ReactomeGSA R package and PADOG.

### Example scRNA-seq Analysis

Raw read counts of the scRNA-seq dataset by Jerby-Arnon *et al.*^*28*^ were retrieved from the Gene Expression Omnibus (GEO, identifier GSE115978). The data was processed using Seurat version 3.1 ^15^ following the new scTransform normalisation strategy regressing out the patient and cohort properties. In order to identify the B cells from the total number of cells we used the first 35 components of the principal component analysis for the subsequent steps. The neighbour graph and clustering was performed using the default parameters. B cell clusters were identified based on a high expression of CD20 (MS4A1), CD79A, CD19, and CD138 (SDC1).

B cells were extracted from the dataset and re-processed, starting with the normalisation step. Here, the top 11 components of the principal component analysis were used for the respective analysis steps. B cell clusters were subsequently classified following the strategy by Sanz *et al.*^*29*^. Plasmablast-like B cells and plasma cells were differentiated based on a low expression of MS4A1 (CD20) in plasmablast-like B cells. Finally, the ssGSEA analysis was performed using the ReactomeGSA R packages’ analyse_sc_clusters function.

The complete workflow including the detailed versions of all used R packages can be found in the respective Jupyter notebook (see Data availability).

### Data Availability

The complete source code of the ReactomeGSA backend, the web-based pathway browser, and the ReactomeGSA Bioconductor R package are available under a permissive open source license on GitHub (https://github.com/reactome). All docker images of the ReactomeGSA analysis system are publically available on Docker Hub (https://hub.docker.com). Central links to all components of the ReactomeGSA system can be found at https://reactome.github.io/ReactomeGSA. The source code of the backend (ie. the Kubernetes application) can be found at https://github.com/reactome/gsa-backend. The source code of the R package is available at https://github,com/reactome/ReactomeGSA. Additionally, a detailed documentation on how to set up the ReactomeGSA analysis system on a local Kubernetes instance can be found on https://reactome.github.io/ReactomeGSA.

The detailed API specification of the ReactomeGSA system is available on https://gsa.reactome.org. Therefore, the complete analysis capabilities can easily be integrated into any other existing software platform.

The code to analyse the example datasets presented in this manuscript can be found as Jupyter notebooks on https://github.com/Reactome/ReactomeGSA-tutorials.

## Acknowledgements

This project received funding from the Research Executive Agency (REA) under the European Union’s Horizon 2020 research and innovation programme under Grant Agreement No. 788042, the US National Institutes of Health (P41 HG003751), and the European Molecular Biology Laboratory. This work was further supported by the FWF-Austrian Science Fund (project P30325-B28).

## Author Contributions

**Hypothesis, study protocol & funding** JG, HHe, **Backend & R package** JG, **Pathway browser** GV, KS, AF, **Example data** JG, VN, **Manuscript** JG, GV, VN, HHe All authors contributed to the final version of the manuscript.

## References

1. The Gene Ontology Consortium. The Gene Ontology Resource: 20 years and still GOing strong. Nucleic Acids Res. 47, D330–D338 (2019).

2. Kanehisa, M., Furumichi, M., Tanabe, M., Sato, Y. & Morishima, K. KEGG: new perspectives on genomes, pathways, diseases and drugs. Nucleic Acids Res. 45, D353–D361 (2017).

3. Subramanian, A. et al. Gene set enrichment analysis: a knowledge-based approach for interpreting genome-wide expression profiles. Proc. Natl. Acad. Sci. U. S. A. 102, 15545–15550 (2005).

4. Jassal, B. et al. The reactome pathway knowledgebase. Nucleic Acids Res. 48, D498–D503 (2020).

5. Mi, H. et al. PANTHER version 11: expanded annotation data from Gene Ontology and Reactome pathways, and data analysis tool enhancements. Nucleic Acids Res. 45, D183–D189 (2017).

6. Huang, D. W., Sherman, B. T. & Lempicki, R. A. Systematic and integrative analysis of large gene lists using DAVID bioinformatics resources. Nat. Protoc. 4, 44–57 (2009).

7. Fabregat, A. et al. Reactome pathway analysis: a high-performance in-memory approach. BMC Bioinformatics 18, 142 (2017).

8. Pilarczyk, M. et al. Connecting omics signatures of diseases, drugs, and mechanisms of actions with iLINCS. Bioinformatics 13 (2019).

9. Papatheodorou, I. et al. Expression Atlas update: from tissues to single cells. Nucleic Acids Res. 48, D77–D83 (2020).

10. Tarca, A. L., Draghici, S., Bhatti, G. & Romero, R. Down-weighting overlapping genes improves gene set analysis. BMC Bioinformatics 13, 136 (2012).

11. Ritchie, M. E. et al. limma powers differential expression analyses for RNA-sequencing and microarray studies. Nucleic Acids Res. 43, e47 (2015).

12. Barbie, D. A. et al. Systematic RNA interference reveals that oncogenic KRAS-driven cancers require TBK1. Nature 462, 108–112 (2009).

13. Hänzelmann, S., Castelo, R. & Guinney, J. GSVA: gene set variation analysis for microarray and RNA-seq data. BMC Bioinformatics 14, 7 (2013).

14. McCarthy, D. J., Chen, Y. & Smyth, G. K. Differential expression analysis of multifactor RNA-Seq experiments with respect to biological variation. Nucleic Acids Res. 40, 4288–4297 (2012).

15. Stuart, T. et al. Comprehensive Integration of Single-Cell Data. Cell 177, 1888–1902.e21 (2019).

16. Amezquita, R. A. et al. Orchestrating single-cell analysis with Bioconductor. Nat. Methods 17, 137–145 (2020).

17. Griss, J. et al. B cells sustain inflammation and predict response to immune checkpoint blockade in human melanoma. Nat. Commun. 10, 4186 (2019).

18. Cancer Genome Atlas Network. Genomic Classification of Cutaneous Melanoma. Cell 161, 1681–1696 (2015).

19. Cancer Genome Atlas Research Network. Comprehensive molecular profiling of lung adenocarcinoma. Nature 511, 543–550 (2014).

20. Cancer Genome Atlas Research Network. Comprehensive genomic characterization of squamous cell lung cancers. Nature 489, 519–525 (2012).

21. Cancer Genome Atlas Research Network. Integrated genomic analyses of ovarian carcinoma. Nature 474, 609–615 (2011).

22. Cancer Genome Atlas Network. Comprehensive molecular portraits of human breast tumours. Nature 490, 61–70 (2012).

23. Mertins, P. et al. Proteogenomics connects somatic mutations to signalling in breast cancer. Nature 534, 55–62 (2016).

24. Zhang, H. et al. Integrated Proteogenomic Characterization of Human High-Grade Serous Ovarian Cancer. Cell 166, 755–765 (2016).

25. Cabrita, R. et al. Tertiary lymphoid structures improve immunotherapy and survival in melanoma. Nature 577, 561–565 (2020).

26. Helmink, B. A. et al. B cells and tertiary lymphoid structures promote immunotherapy response. Nature 577, 549–555 (2020).

27. Lu, Y. et al. Complement Signals Determine Opposite Effects of B Cells in Chemotherapy-Induced Immunity. Cell 180, 1081–1097.e24 (2020).

28. Jerby-Arnon, L. et al. A Cancer Cell Program Promotes T Cell Exclusion and Resistance to Checkpoint Blockade. Cell 175, 984–997.e24 (2018).

29. Sanz, I. et al. Challenges and Opportunities for Consistent Classification of Human B Cell and Plasma Cell Populations. Front. Immunol. 10, 2458 (2019).

30. Pinto, D. et al. A functional BCR in human IgA and IgM plasma cells. Blood 121, 4110–4114 (2013).

31. Gromnitza, S. et al. Tropomyosin-related kinase C (TrkC) enhances podocyte migration by ERK-mediated WAVE2 activation. FASEB J. 32, 1665–1676 (2018).

32. Kurebayashi, Y. et al. Comprehensive Immune Profiling of Lung Adenocarcinomas Reveals Four Immunosubtypes with Plasma Cell Subtype a Negative Indicator. Cancer Immunol Res 4, 234–247 (2016).

33. Tarca, A. L., Bhatti, G. & Romero, R. A comparison of gene set analysis methods in terms of sensitivity, prioritization and specificity. PLoS One 8, e79217 (2013).

34. MacNeil, S. M., Johnson, W. E., Li, D. Y., Piccolo, S. R. & Bild, A. H. Inferring pathway dysregulation in cancers from multiple types of omic data. Genome Med. 7, 61 (2015).

35. Vaske, C. J. et al. Inference of patient-specific pathway activities from multi-dimensional cancer genomics data using PARADIGM. Bioinformatics 26, i237–45 (2010).

36. Sedgewick, A. J., Benz, S. C., Rabizadeh, S., Soon-Shiong, P. & Vaske, C. J. Learning subgroup-specific regulatory interactions and regulator independence with PARADIGM. Bioinformatics 29, i62–70 (2013).

37. Huber, W. et al. Orchestrating high-throughput genomic analysis with Bioconductor. Nat. Methods 12, 115–121 (2015).

38. UniProt Consortium. UniProt: a worldwide hub of protein knowledge. Nucleic Acids Res. 47, D506–D515 (2019).

39. Orchard, S. et al. The MIntAct project--IntAct as a common curation platform for 11 molecular interaction databases. Nucleic Acids Res. 42, D358–63 (2014).

40. Law, C. W., Chen, Y., Shi, W. & Smyth, G. K. voom: Precision weights unlock linear model analysis tools for RNA-seq read counts. Genome Biol. 15, R29 (2014).

41. McCarthy, D. J., Campbell, K. R., Lun, A. T. L. & Wills, Q. F. Scater: pre-processing, quality control, normalization and visualization of single-cell RNA-seq data in R. Bioinformatics 33, 1179–1186 (2017).

42. Colaprico, A. et al. TCGAbiolinks: an R/Bioconductor package for integrative analysis of TCGA data. Nucleic Acids Res. 44, e71 (2016).

